# A simple high-throughput approach identifies actionable drug sensitivities in patient-derived tumor organoids

**DOI:** 10.1101/138412

**Authors:** Nhan Phan, Jenny J. Hong, Bobby Tofig, Matthew Mapua, David Elashoff, Neda A. Moatamed, Jin Huang, Sanaz Memarzadeh, Robert Damoiseaux, Alice Soragni

## Abstract

There is increasing interest in developing 3D tumor organoid models for drug development and personalized medicine applications. While tumor organoids are in principle amenable to high-throughput drug screenings, progress has been hampered by technical constraints and extensive manipulations required by current methodologies. Here, we introduce a miniaturized, fully automatable, flexible high-throughput method using a simplified geometry to rapidly establish 3D organoids from cell lines and primary tissue and robustly assay drug responses. As proof of principle, we use our miniring approach to establish organoids of high-grade serous tumors and one carcinosarcoma of the ovaries and screen hundreds of protein kinase compounds currently FDA-approved or in clinical development. In all cases we could identify drugs causing significant reduction in cell viability, number and size of organoids within a week from surgery, a timeline compatible with therapeutic decision making.

## Introduction

Cancer therapy is rapidly progressing toward individualized regimens not based on the organ of origin, but rather on the molecular characteristics of tumors. Next generation sequencing (NGS) is typically regarded as the key to access this potentially actionable molecular information^1,2^. However, recent studies showed how only a small number of cancers can be singled out and targeted with this approach, in part because very few gene alteration-drug pairs are unequivocally established and few accurate predictive biomarkers are available^3–7^. Thus, functional precision therapy approaches where the primary tumor tissue is directly exposed to drugs to determine which may be efficacious have the potential to boost personalized medicine efforts and influence clinical decisions^3,4^. Establishing patient-derived xenografts (PDX) is a costly and time consuming option that only allows to screen very few potential drugs. Conversely, ex vivo 3D tumor spheroids or organoids derived from primary cancers can be easily established and potentially scaled to screen hundreds to thousands of different conditions.

3D cancer models have been consistently shown to faithfully recapitulate features of the tumor of origin in terms of cell differentiation, heterogeneity, histoarchitecture and clinical drug response^4,8–15^. Various methods to set up tumor spheroids or organoids have been proposed, including using low-attachment U-bottom plates, feeding layers or various biological and artificial matrices^9,12,13,16–22^. Methods using low-attachment U-bottom plates ideally only carry one organoid per well, have limited automation and final assay capabilities^18–20^. In addition, not all cells are capable of forming organized 3D structures with this method. Approaches that include a bio-matrix, such as Matrigel, have the potential to offer a scalable alternative in which cancer cells thrive^9,14,23,24^. However, most approaches so far rely on thick volumes of matrix which is not cost-effective, potentially hard for drugs to efficiently penetrate and difficult to dissolve fully at the end of the experiment^23^. In other applications, organoids are first formed and then transferred to different plates for drug treatment or final readout which can result in the tumor spheres sticking to plastic or breaking^14,24^. In addition, some assays require to disrupt the organoids to single cell suspensions at the end of the experiment^16,22^. All of these manipulations introduce significant variability thus limiting applicability in screening efforts^12^.

To overcome these limitations, we introduce a facile assay system to screen 3D tumor organoids that takes advantage of a specific geometry. Our miniaturized ring methodology does not require functionalized plates. Organoids are assayed in the same plate where they are seeded, with no need for sample transfer at any stage or dissociation of the pre-formed tumor organoids to single cell suspensions. Here, we show that the mini-ring approach is simple, robust, requires few cells and can be easily automated for high-throughput applications. Using this method, we were able to rapidly identify clinically actionable drug sensitivities for several ovarian cancers and high-grade serous tumors by testing two different drug concentrations and a library of 240 protein kinase inhibitor compounds.

## Results

### Establishment of 3D tumor models in ring format

In order to rapidly screen organoids, we first established a miniaturized system that allows to setup hundreds of wells and perform assays with minimal manipulation. We adapted the geometry used to plate tumor cells in Matrigel to generate mini-rings around the rim of the wells. This is attained by plating single cell suspensions obtained from a cell line or a surgical specimen pre-mixed with cold Matrigel (3:4 ratio) in a ring shape around the rim in 96 well plates (Fig. 1a). Rings can be established using a single-well or multichannel pipette. Use of a robotic system or automated 96 well pipettor is theoretically feasible as long as temperature and plate positioning can be effectively controlled. The combination of small volume plated (10 *µ*l) and surface tension holds the cells in place until the Matrigel solidifies upon incubation at 37°C and prevents 2D growth at the center of the wells. The ring configuration allows for media addition and removal so that changes of conditions or treatment addition to be easily performed by pipetting directly in the center of the well, preventing any disruption of the gel.

**Figure 1.**
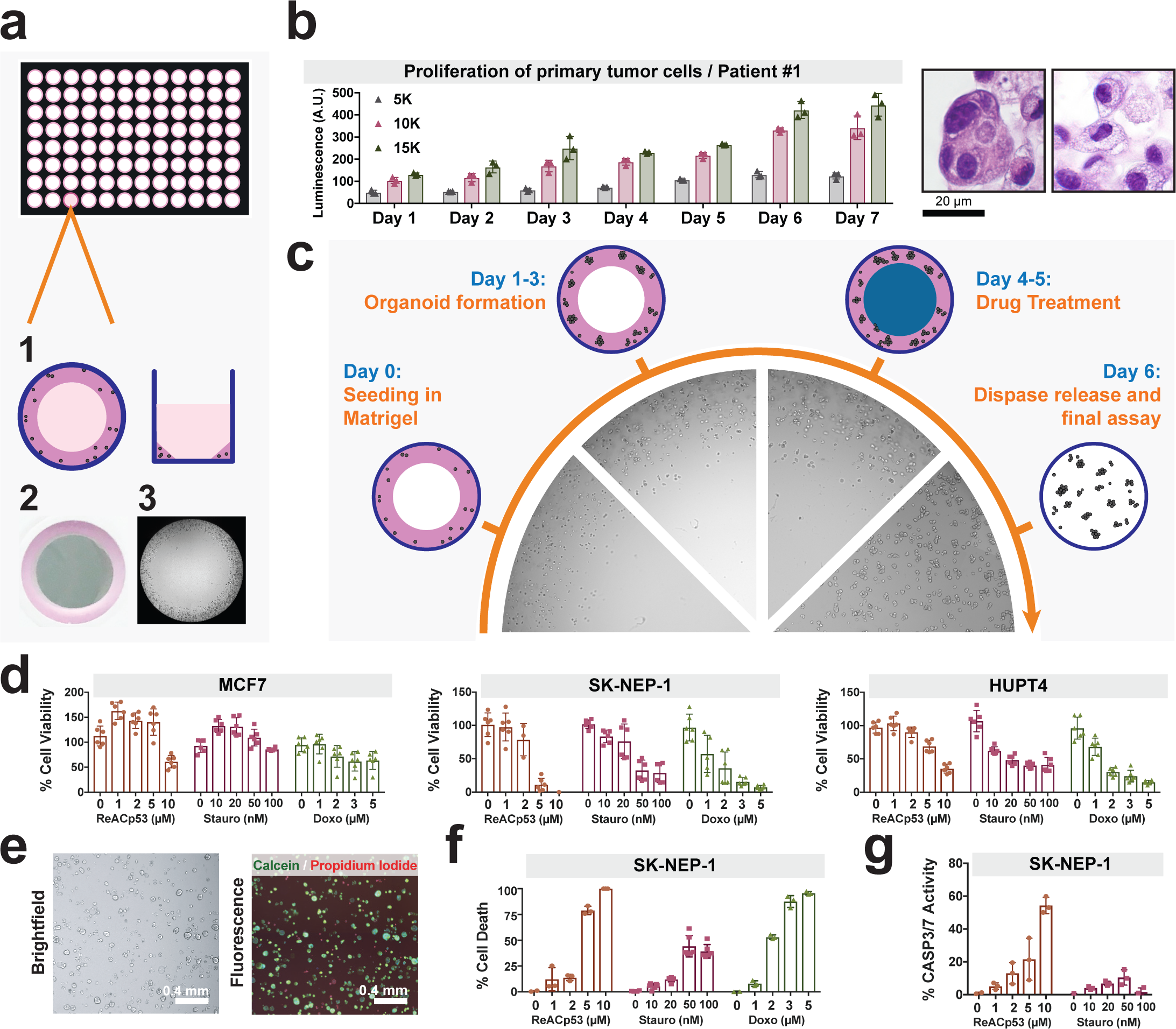
The mini-ring method for 3D tumor cell biology. (a) Schematics of the mini-ring setup. Cells are plated in Matrigel around the rim of the wells form a solid thin ring as depicted in 1 and photographed in 2, which has decreasing thickness. The picture in 3 acquired with a cell imager shows tumor organoids growing at the periphery of the well as desired, with no invasion of the center. (b) Proliferation of primary tumor cells as measured by ATP release. Different seeding densities were tested and compared (5, 10 and 15K cells). The mini-ring method allowed the patient sample to grow and maintain the heterogeneity and histology of the original ovarian tumor which had a high-grade serous carcinoma component (H&E left picture) and a clear cell component (H&E right picture). (c) Schematic of the drug-treatment experiments performed in the mini-ring setting. The pictures are representative images as acquired using a Celigo cell imager. (d - g) Assays to monitor drug response of cell lines using the mini-ring configuration. Three drugs (ReACp53, Staurosporine and Doxorubicin) were tested at five concentrations in triplicates for all cell lines. (d) ATP release assay (CellTiter-Glo 3D) readout. (e) and (f) Calcein/PI readout. (e) Representative image showing staining of MCF7 cells with the dyes and segmentation to quantify the different populations (live / dead). (f) Quantification of Calcein/PI assay for three-drug assay. (g) Quantification of cleaved caspase 3/7 assay. Doxorubicin was omitted due to its fluorescence overlapping with the caspase signal. For all graphs, symbols are individual replicates, bars represent the average and error bars show SD.

Cancer cell lines grown in mini-ring format give rise to organized tumor organoids that recapitulate features of the original histology (Fig. S1 and Table S1).

Similarly, we can routinely establish patient-derived tumor organoids (PDTOs) using the same geometry. As an example, Patient #1 was diagnosed with a high-grade mixed type carcinoma with both a serous component as well as a clear cell component (Table S1 and Fig. S2). Similarly, cells isolated from Patient #1 tumor grown in our ring system show two distinct cytomorphologies: one group of cells have clear cytoplasm and cuboidal appearance while the second group of cells organize in clusters in a columnar fashion and have dense cytoplasm (Fig. S2). These morphologies are compatible with the two different histologies found in the original tumor, clear cell and high-grade serous carcinoma (Fig. S2a). Moreover, both the tumor organoids and the primary cancer show similar p53 staining patters, with populations of p53-positive and p53-negative cells (Fig. S2b-c). Thus, patient samples obtained at the time of surgery can proliferate in our system and maintain the heterogeneity of the original tumor as expected (Fig. 1b and S2).

### Assay optimization

Next, we optimized treatment protocols and readouts for the mini-ring approach. Our standardized paradigm includes: seeding cells on day 0, establishing organoids for 2-3 days followed by two consecutive daily drug treatments, each performed by complete medium change (Fig. 1c). To demonstrate feasibility, we performed small scale screenings testing three drugs at five different concentrations in triplicates, ReACp53^16^, Staurosporine^25^ and Doxorubicin (Fig. 1d-g). We optimized different readouts in order to adapt the method to a specific research question or instrument availability. After seeding cells in standard white plates, we performed a luminescence-based ATP assay to obtain a metabolic readout of cell status, calculate EC_50_s ranged between 2.5 *µ*M (MDA-MB-468) and 10 *µ*M (MCF7) for ReACp53, between 100 nM (MCF7) and 800 nM (PANC 03.27) for Staurosporine and between 900 nM (SKNEP) and 12 *µ*M (MCF7) for Doxorubicin. For instance, our measurements are in line with the Doxorubicin resistance of MCF7 cells grown in Matrigel in 3D that has been previously reported^26^.

We performed two consecutive treatments which allows the drugs to not only penetrate the gel but also to reach organoids that may be bulky^16^. However, the assay is flexible and can be easily adapted to single treatments followed by longer incubations, multiple consecutive recurring treatments, multi-drug combinations or other screening strategies (Fig. S4).

We also implemented assays to quantify drug response by measuring cell viability after staining of live organoids with specific dyes followed by imaging. We optimized a calcein-release assay coupled to propidium iodide (PI) staining as well as a caspase 3/7 cleavage assay that can be readily performed after seeding the cells in standard black plates (Fig. 1e-g and S5). For all assays, tumor organoids are stained following dispase release. After a 40-minute incubation, organoids are imaged, and pictures are segmented and quantified (Fig. 1e-g and S4). All the assays are performed within the same well in which spheroids are seeded. Although the various assays we introduce are testing different aspects of cell viability and measure distinct biological events, results were mostly concordant across the methods for the three drugs tested (Fig. 1 and S5).

### Identification of actionable drug responses in PDTOs

A rapid functional assay to determine drug sensitivities of primary specimens can offer actionable information to help tailoring therapy to individual cancer patients^3^. We tested suitability of our approach to rapidly and effectively identify drug susceptibilities of three ovarian cancer samples and one high-grade serous peritoneal cancer specimen obtained from the operating room (Table S1; Fig. 2 and 3). In all cases, ascites or tumor samples were processed after surgery (see Methods) and then plated as mini-rings as described above. In order to maximize the amount of information extracted from irreplaceable clinical samples, we investigated the possibility to concurrently perform multiple assays on the same plate. To do so, we first optimized the initial seeding cell number (5000 cells/well) to couple an ATP metabolic assay to 3D tumor count and total organoid area measurements. This seeding density yields a low-enough number of organoids to facilitate size distribution analysis but sufficient ATP signal to be within the dynamic range of the CaspaseGlo 3D assay. For each patient sample, we seeded six 96 well plates and tested 240 protein kinase inhibitors FDA-approved or in clinical development. We tested each drug at two different concentrations (120 nM and 1 *µ*M), for a total of 480 different conditions tested. Differently from established cancer cell lines, the number of cells obtained from surgical specimens can be limiting. As such, a one- or two-dose primary screening is a reasonable and commonly adopted approach to identify potential hits. Validation can then be performed using frozen aliquots of cells that we cryopreserve after tissue processing post-surgery.

**Figure 2.**
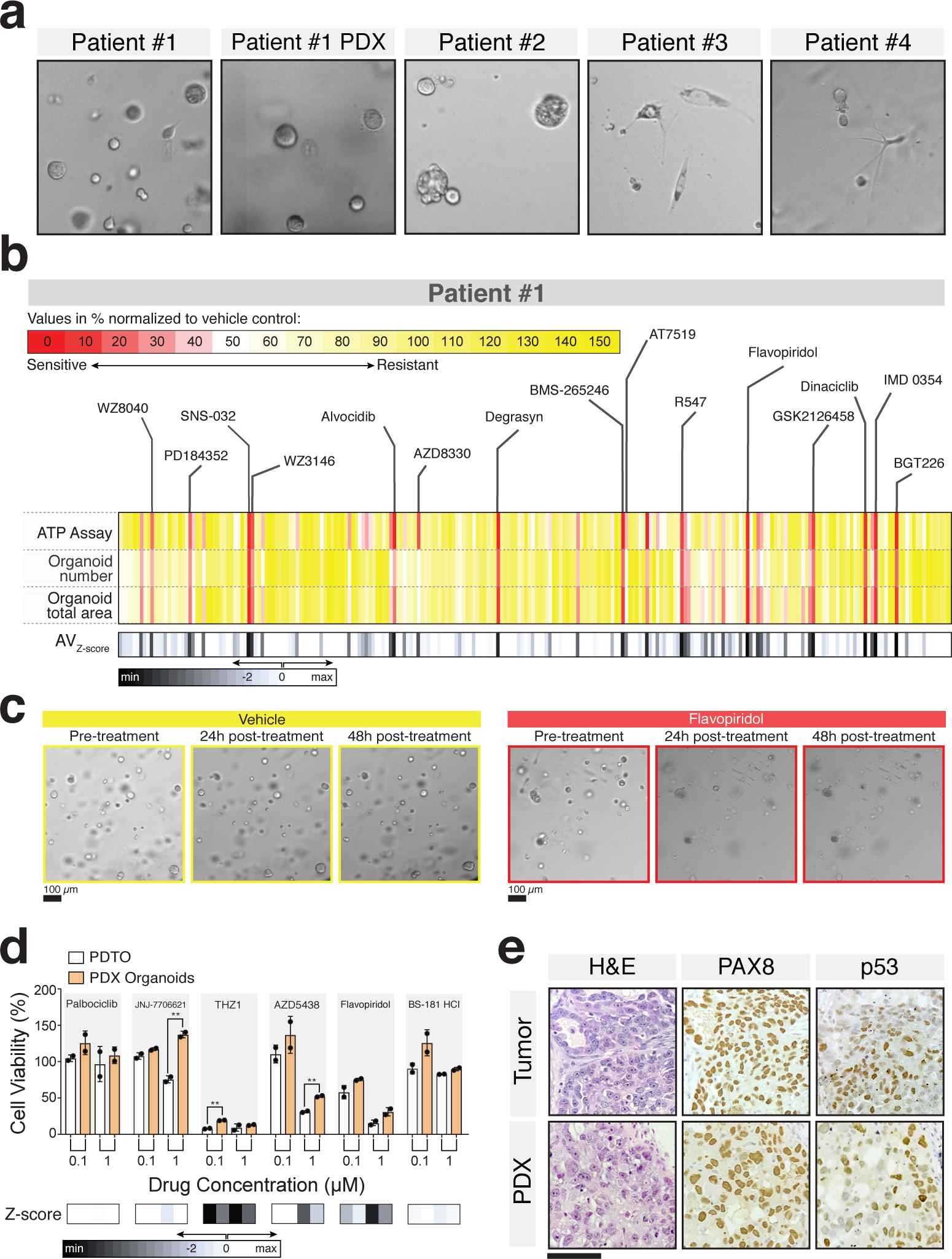
Mini-ring approach to unveil drug response patterns in PDTOs. **(a)** Morphology of all PDTOs established in this study as visualized by brightfield microscopy. **(b)** Results of kinase screening experiment for Patient #1 PDTOs. Three readouts were used for this assay: ATP quantification as measured by CellTiter-Glo 3D and organoid number or size quantification evaluated by brightfield imaging. Brightfield images were segmented and quantified using the Celigo S Imaging Cell Cytometer Software. Both organoid number as well as total area were evaluated for their ability to capture response to drugs. In this plot, each vertical line is one drug, all 240 tested are shown. Values are normalized to the respective vehicle controls for each method and expressed as %. Average_Z-score_ calculated as reported in Methods. **(c)** A representative image of the effects of the indicated drug treatments as visualized by the Celigo cell imager. **(d)** Small scale kinase assay on Patient #1 primary PDTOs and PDX-derived cells. ATP readout. Four molecules not present in the primary screening were tested. Flavopiridol and BS-181 HCl are included as positive and negative control respectively. **(e)** Comparison of the histology of the primary tumor with the established PDX. Scale bar: 100 *μ*m.

**Figure 3.**
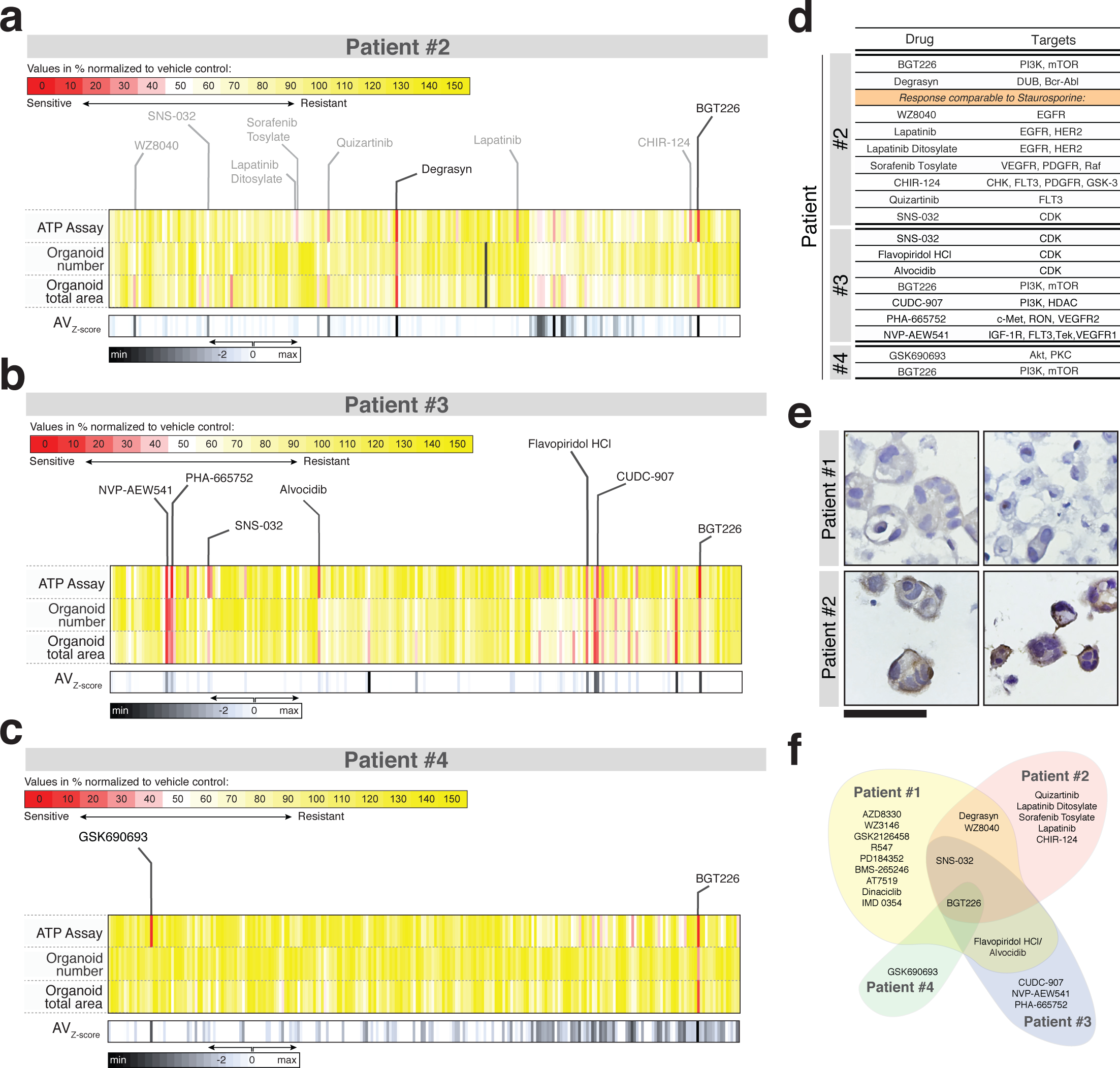
Individualized response of PDTOs to tyrosine kinase inhibitors. **(a), (b)** and **(c)** Results of kinase screening experiment on Patients 2-4 organoids. Each vertical line represents one of 241 tested drugs. Values are normalized to the respective vehicle controls (DMSO) for each method and expressed as %. **(d)** Table of drug leads causing ~75% cell death. For Patient #2, we included drugs inducing a response comparable to the Staurosporine control (~60% cell death). **(e)** Expression of the multi-drug efflux protein ABCB1 in PDTOs as visualized by IHC. Patient #2 expresses very high levels of the ABC transporter. Scale bar: 60 *μ*m. **(f)** Diagram illustrating limited overlap between the detected patterns of response identified through the mini-ring assay for all patients.

For PDTOs, we used the same experimental paradigm optimized above. All steps (media change, drug treatment) were automated and performed in less than 2 minutes/plate using a Beckman Coulter Biomek FX integrated into a Thermo Spinnaker robotic system. At the end of each experiment, PDTOs are first imaged in brightfield mode for organoid count/size distribution analysis followed by an ATP assay performed on the same plates. The measurements yielded high quality data that converged on several hits, highlighting the feasibility of our approach to identify potential leads (Fig. 2, 3 and S7).

### Patient #1: High-grade mixed type carcinoma

Cells obtained from Patient #1 at the time of cytoreductive surgery were chemo-naïve, and the heterogeneous nature of this clear cell/HGSC tumor was recapitulated in the PDTOs (Table 1, Fig. 1b and S2). Despite aggressive debulking surgery and treatment with carboplatin and paclitaxel regimens, Patient #1 had persistent disease, never achieved complete remission and overall survival from diagnosis was 11 months. Resistance to carboplatin was also observed in our high-throughput assay, with no significant reduction of viability observed at either 10 or 25 *µ*M concentrations (Fig. S6). The organoids were however sensitive to ~6% of the protein kinase inhibitors tested (16/240), with sensitivity defined as residual cell viability ≤ 25% and average Z-score ≤ −5 (Fig. 2b and Table 1; see Methods for Z-score calculations). Patient #1’s tumor organoids responded to 58% of all cyclin-dependent kinase (CDK) inhibitors tested (7/12 total, 11 different compounds and one, Flavopiridol, in two formulations). In particular, cells appeared highly sensitive to inhibitors hitting CDK1/2 in combination with CDK 4/6 or CDK 5/9 (Table 1, Fig. 2c and S7a). Interestingly, CDK inhibitors have found limited applicability in ovarian cancer therapy so far^27^. Based on the profiles of the CDK inhibitors tested and on the response observed (Fig. S7a), we selected four untested molecules to assay. We anticipated that Patient #1 would not respond to Palbociclib (targeting only CDK4/6) and THZ1 (CDK7) while expecting a response to JNJ-7706621 (CDK1/2/3/4/6) and AZD54338 (CDK1/2/9; Fig. S7a). BS-181 HCl and AZD54338 (CDK1/2/9; Fig. S7a). BS-181 HCl and Flavopiridol were included as negative and positive control respectively. Results show that organoids were not sensitive to JNJ-7706621 but had a strong response to THZ1 (Fig. 2d). Both THZ1 and BS-181 HCl specifically target CDK7. Nevertheless, Patient #1 PDTOs showed a strong response to the former but no response to the latter which could be attributed to the different activity of the two as recently observed in breast cancer^28^.

**Table 1.** List of molecules causing >75% reduction in viability in PDTOs established from Patient #1’s tumor.

| Patient #1 | Drug | Target | AV<sub>Z-score</sub> | Viability % |
| --- | --- | --- | --- | --- |
|  | SNS-032 | CDK | -8.0 | 6.7 |
|  | Alvocidib | CDK | -7.7 | 2.9 |
|  | AT7519 | CDK | -7.3 | 4.2 |
|  | BMS-265246 | CDK | -7.6 | 5.0 |
|  | Flavopiridol HCl | CDK | -8.6 | 6.0 |
|  | R547 | CDK | -8.1 | 11.8 |
|  | Dinaciclib | CDK | -8.3 | 8.1 |
|  | Degrasyn | DUB, Bcr-Abl | -7.6 | 5.4 |
|  | WZ3146 | EGFR | -6.9 | 16.2 |
|  | WZ8040 | EGFR | -6.5 | 22.0 |
|  | IMD 0354 | IKK | -8.6 | 4.8 |
|  | PD184352 | MEK | -6.7 | 20.1 |
|  | AZD8330 | MEK | -6.1 | 15.5 |
|  | GSK2126458 | PI3K, mTOR | -7.9 | 11.6 |
|  | BGT226 | PI3K, mTOR | -8.5 | 5.0 |

We also attempted to validate the screening results in vivo establishing patient-derived xenografts (PDX) by injecting Patient #1 cells subcutaneously in NSG mice (500K/mouse, 12 mice). However, only three mice developed PDXs over the course of five months. The xenografts resembled the histology of the primary tumor (Fig. 2e). In order to test whether the PDXs had a similar response to CDK inhibitors, we dissociated the PDX to single cell suspension and generated organoids from one of them (Fig. 2a, d-e). The PDX-derived organoids showed an overall trend toward a reduction in sensitivity to CDKs when compared to the PDTOs. We observed a statistically significant decrease in response to 0.1 *μ*M THZ1, and 1 *μ*M JNJ-7706621 and AZD5438 (p<0.01, Fig. 2d) in the PDX-derived organoid compared to the PDTOs. This is not unexpected, as human cancer cells grown in mice rapidly diverge from the tumor they were obtained from^29,30^.

### Patient #2: Platinum-resistant high-grade serous ovarian carcinoma

Patient #2 was diagnosed with progressive, platinum-resistant high-grade serous ovarian cancer (HGSOC) and was heavily pretreated prior to sample procurement (Table S1). Patient #2 PDTOs showed a strong response (residual cell viability ≤ 25% and average Z-score ≤ −5) to only 0.8% of all drugs tested (2/240, Fig. 3a, 3d and S7b). PDTOs were also platinum-resistant in our system (Fig. S6), with no reduction of viability observed upon treating the cells with either 10 or 25 *µ*M carboplatin.

Remarkably, Patient #2 PDTOs showed a moderate response to our positive control, Staurosporine, a pan-kinase inhibitor with very broad activity^25^. The significant lack of response to multiple therapies observed for Patient #2 led us to hypothesize that there could be over-expression of efflux membrane proteins. Indeed, the PDTOs showed a high level of expression of ABCB1 (Fig. 3e). High expression of the ATP-dependent detox protein ABCB1 is frequently found in chemoresistant ovarian cancer cells and recurrent ovarian cancer patients’ samples and has been correlated with poor prognosis^31,32^.

Flavopiridol were included as negative and positive control respectively. Results show that organoids were not sensitive to JNJ-7706621 but had a strong response to THZ1 (Fig. 2d). Both THZ1 and BS-181 HCl specifically target CDK7. Nevertheless, Patient #1 PDTOs showed a strong response to the former but no response to the latter which could be attributed to the different activity of the two as recently observed in breast cancer^28^.

We found a moderate response, comparable to the effect of Staurosporine (~40% residual cell viability), to EGFR or EGFR/HER2 inhibitors including Lapatinib and WZ8040 (Fig. 3d). We could detect high expression of EGFR at the plasma membrane of the tumor cells (Fig. S7e), as is common for platinum-resistant ovarian cancer^33^.

### Patients #3: Carcinosarcoma of the ovary

Patient #3 presented with a carcinosarcoma of the ovary, an extremely rare and aggressive ovarian tumor which has not been fully characterized at the molecular level yet^34,35^ (Table S1, Fig. 3b, 3d and S7c). In our screening, the PDTOs established from this tumor responded to ~3% of all tested kinase inhibitors (7/240, residual cell viability ≤ 25% and average Z-score ≤ −1.5), including CDK inhibitors and PI3K inhibitors.

### Patient #4: High-grade peritoneal carcinoma

Patient #4 was diagnosed with a high-grade peritoneal tumor and showed a response to 0.8% of all tested drugs (2/240, Table S1, Fig. 3c-d and S7d). The PDTOs showed a marked response to two drugs, one pan-Akt inhibitor (GSK690693) and a PI3K/mTOR inhibitor (BGT226), with a measured cell viability ≤ 25% and average Z-score ≤ −5. However, differently from Patient #2, Patient #4 PDTOs were sensitive to Staurosporine, with only 9±1% residual viability after 2 days of treatment. Protein kinase C (PKC), which is the primary target of Staurosporine, is also a secondary target of GSK690693^36^.

Although only 2 inhibitors caused a 75% reduction in cell viability, 11 agents caused ≥ 50% cell death (Z-score ≤ −5). Using this cutoff, we could identify 6 mTOR inhibitors including Omipalisib, Apitolisib, and Sapanisertib. These constitute 30% of all the mTOR inhibitors tested, pinpointing a potential vulnerability of the pathway for this PDTO sample.

## Discussion

We devised and optimized a facile high-throughput approach to establish and screen 3D models and tumor organoids established from cell lines or clinical samples. We used our approach to functionally profile four tumors obtained from surgeries. We observed highly tumor-specific responses, with very little overlap among the inhibitors that each clinical sample was sensitive to. Only one molecule, BGT226, showed activity in all tumors (Fig. 3f). A phase I basket trial of this PI3K/mTOR inhibitor showed moderate responses in unstratified patients^37^. In this trial as in trials of other PI3K/mTOR inhibitors, there are no markers to clearly and unambiguously identify patients that will benefit from therapy^38,39^. PI3K/mTOR inhibitors are just one example of drugs for which a clear predictive biomarker is lacking. In fact, the absence of specific, unequivocal biomarkers predictive of response is a common limitation and challenge associated with many kinase inhibitors^40^. Our assay could bypass the lack of biomarkers, and identify sensitive cases from a functional standpoint. Thus, patients may greatly benefit from functional PDTO testing, either to identify a suitable therapy or to facilitate patient selection for clinical trials^3,4,12,14,41^.

A recent study by Vlachogiannis et al found that patient-derived organoids could accurately predict patient responses to therapy, with 100% sensitivity and 93% specificity^4^. In our experiments, we could recapitulate the carboplatin resistance of patients ex vivo (Fig. S6). Interestingly, PDTOs exhibited differential responses to different molecules targeting the same pathway (Fig. 2–3 and S7). As an example, CDKs were obvious sensitive targets for inhibition in Patient #1 PDTOs (Table 1). However, when we attempted to use the information collected from the screening to identify additional CDK inhibitors with similar target profiles that would elicit expected responses, we were only partially successful (Fig. 2d and S7a). This could be due to different efficacies^28^, secondary targets or other properties of the inhibitors. Therefore, our high-throughput approach allows not only to identify susceptible pathways, but also to select the most effective agent from a class of molecules.

One important advantage of the mini-ring approach is the small number of cells needed. This allows testing samples as obtained from biopsies or surgeries without the need for expansion in vitro or in vivo, a process which can lead to substantial divergence from the tumor of origin^29,42^. In our experience, the vast majority of solid tumor specimens do not adhere or grow in 2D, which limits the possibility of expansion in vitro. Moreover, take rates of patient-derived tumor cells in vivo can be highly variable^43^. We could only generate a limited number of PDXs from 6 millions of Patient #1 cells over five months (3/12), while we could test 240 drugs in five days with a fifth of the cells. Therefore, our approach can be very effective to test patient samples that are recalcitrant to grow in vivo, reducing times and costs (Fig. 2d-e).

Another interesting application of PDTO screenings for precision medicine applications is in the rare disease space. We could find several effective molecules against a carcinosarcoma of the ovary (Fig. 3b and 3d). The rarity of this type of cancer, which accounts for only 1-4% of all ovarian tumors^35^, hinders the design of clinical trials to identify effective regimens, and therapy is often modeled on other cancer types. For instance, a clinical trial that demonstrated the efficacy of platinum-agents in this setting run for approximately 20 years to enroll 136 patients^44^. The ability to model rare tumors using PDTOs and perform robust screenings ex vivo offers an opportunity to identify drugs in a disease- and mechanism-agnostic manner, even for tumor types that are largely uncharacterized.

## Conclusions

High-throughput drug screenings using PDTOs have many advantages and a real opportunity to be factored into therapeutic decisions. Our methodology can be a robust tool to standardize functional precision medicine efforts^3^, given its ease of applicability to many different systems and drug screening protocols (Fig. 1, S3 and S4). While we used the mini-ring setup for drug screening purposes, the same methodology is suitable for studies aimed at characterizing organoids’ biological and functional properties with medium-to high-throughput. Complete automation, scalability to 384 well plates, and flexibility to use different supports beside Matrigel can further facilitate broader implementation of the mini-ring approach.

## Acknowledgments

This project was supported by a Worldwide Cancer Research grant to AS (#16-0253). We acknowledge support by the Hirshberg Foundation (to AS), the NIH/ National Center for Advancing Translational Science UCLA CTSI Grant UL1TR000124 (to AS), the UCLA SPORE in Prostate Cancer (NIH P50CA092131, PI: Robert Reiter), an AACR-Millennium Fellowship in Prostate Cancer Research (#14-40-38-SORA to AS), an American Cancer Society Research Scholar Grant (#RSG-14-217-01-TBG to SM) and a Jonsson Cancer Center Foundation Impact Award (to SM).

## Author Contributions

AS and NP designed the project and carried out the experiments. JJH and MM performed experiments on clinical samples. SM obtained the patient samples. JH contributed to feasibility experiments. BT and RB generated the kinase inhibitor drug library and optimized automation for primary sample assays. NAM analyzed tissue and organoid sections. DE contributed to data analysis. AS analyzed the data and wrote the paper with contributions from all authors.

## Methods

### Cell lines and primary samples

Cell lines are cultured in their recommended medium in the presence of 10% FBS (Life Technologies #10082-147) and 1% Antibiotic-Antimycotic (Gibco). DU145, PC3, PANC1 and HUTP4 were culture in DMEM (Life Technologies #1195-065). PAN03.27, MDA-MB-468 and MCF-7 was cultured in RPMI (Life Technologies #22400-089). SK-NEP-1 was cultured in McCoy medium (ATCC #30-2007). All treatments are performed in serum-free medium (PrEGM, Lonza #CC-3166 or MammoCult, StemCell Technologies # 05620).

### Primary samples

Primary ovarian cancer specimens were dissociated to single cells and cryopreserved or plated right after processing. In short, fresh tumor specimens or ascites samples are obtained from consented patients (UCLA IRB 10-000727). Solid tumor specimens are minced, then enzymatically digested with collagenase IV (200 U/ml). The resulting cell suspension is filtered through a 40 μm cell strainer.

For Patient #1 patient-derived xenografts, 12 NSG mice were injected with 500K cells in matrigel on the flank. Tumor growth was monitored over time. After about 5 months, three mice developed measurable tumors, which were collected at euthanasia. A portion was fixed and processed for histology, and the remaining tissue was dissociated to single cell and assayed as the primary sample from the same patient.

### Chemicals

Doxorubicin hydrochloride was purchased from Sigma (#44583). Staurosporine was purchased from Cell Signaling Technology (#9953S). ReACp53 was synthesized by GL Biochem and prepared as described in Soragni et al, 2016.

### 3D organoids seeding/treatment procedure

Single-cell suspensions (2K-10K/well) were plated around the rim of the well of 96 well plates in a 3:4 mixture of PrEGM medium and Matrigel (BD Bioscience CB-40324). White plates (Corning #3610) were used for ATP assays while black ones (Corning #3603) were used for caspase or calcein assays. Plates are incubated at 37°C with 5% CO_2_ for 15 minutes to solidify the gel before addition of 100 *µ*l of pre-warmed PrEGM or Mammocult to each well using an EpMotion (Eppendorf). Two days after seeding, medium is removed and replaced with fresh PrEGM or Mammocult containing the indicated drugs. The same procedure is repeated daily on two consecutive days. 24h after the last treatments, media is removed and wells are washed with 100 *µ*l of pre-warmed PBS. To prepare for downstream experiments, organoids are then released from Matrigel by incubating for 40 minutes in 50 *µ*l of 5 mg/mL dispase (Life Technologies #17105-041). All steps are performed with the EpMotion for small scale experiments and medium is removed/added from the center of the wells. For the high-throughput kinase screening experiment, we utilized a Beckman Coulter Biomek FX system with 96 channel head integrated into a Thermo Spinnaker robotic system with Momentum scheduling software. In short, an intermediary dilution plate (Axygen P-96-450V-C-S) was filled with 100 *µ*l/ well of media and pre-warmed to 37°C. Using pre-sterilized p50 tips, 1 *µ*l of drug is transferred from a library compound plate to the intermediary media plate and thoroughly mixed. Next, the robot gently removed 100 *µ*l of media from the matrigel/cell plate. The liquid handler was set up to hit the dead center of each well with no contact to the Matrigel mini-ring. As a last step, the robot transferred 100 *µ*l from the intermediary plate (media+drug) to the matrigel/cell plate. Media was easily dispensed without touching or disrupting the Matrigel mini-ring. The total process time outside of the CO_2_ incubator was less than 2 minutes allowing the temperature to be controlled throughout.

### ATP assay

After the organoid release, 75 *µ*l of Celltiter-Glo 3D Reagent (Promega #G968B) is added to each well followed by 1 minute of vigorous shaking. After a 30-minute incubation at room temperature and an additional minute of shaking, luminescence is measured with a SpectraMax iD3 (Molecular Devices) over 500 ms of integration time. Data is normalized to vehicle and plotted and EC_50_ values are calculated with Prism 7.

### High-throughput screening data analysis

For the high-throughput drug screening, DMSO and Staurosporine (1 *µ*M) are used as negative and positive control respectively. Cell viability values are normalized to vehicle (DMSO) and expressed as %. Z-scores are calculated as [(Y_drug X_) - (average Y_vehicle_)]/(average standard deviation_vehicle_), where Y is either viability, organoid total number or organoid area. The average standard deviation_vehicle_ is a single value calculated across all three assay plates to better account for overall variability.

The three Z-scores, one for viability, one for organoid total number and one for organoid area, are then averaged for each drug. This was performed separately for each patient.

Hits are determined following three criteria: (1) cell death shows concentration-dependency, (2) residual cell viability at 1 *µ*M is ≤ 25% and (3) average _Z-score_≤ −5. For Patient #3, an average _Z-score_ cutoff of −1.5 was used. The different threshold was adopted due to heterogeneity in the vehicle standard deviations across subjects.

For Patient #2, partial hits are defined as drugs residual cell viability > 25% and ≤ (Staurosporine + 5%) at 1 *µ*M.

### Caspase 3/7/Hoechst assay

After dispase treatment, 100 *µ*l of Nexcelom ViaStain™ Live Caspase 3/7 staining solution is added to each well. The staining solution consists of 2.5 *µ*M Caspase reagent (Nexcelom #CSK-V0002) and 3 *µ*g/ml Hoechst (Nexcelom #CS1-0128) in serum-free RPMI medium. Plates are incubated 37°C/5% CO_2_ for 45 minutes and imaged with a Celigo S Imaging Cell Cytometer (Nexcelom). Data is normalized to vehicle values and plotted with Prism 7.

### Calcein-AM/Hoechst/Viability assay

For this assay, 100 *µ*l of Calcein-AM/Hoechst/PI viability staining solution are added to each well containing the released organoids. The staining solution includes the Calcein-AM reagent (Nexcelom CS1 #0119; 1:2000 dilution), Propidium Iodide (Nexcelom #CS1-0116; 1:500 dilution), Hoechst (Nexcelom #CS1-0126; 1:2500 dilution) in serum-free RPMI medium. Samples are incubated for 15 minutes at 37°C with 5% CO_2_ before imaging with a Celigo S Imaging Cell Cytometer (Nexcelom).

### Immunohistochemistry

Cells processed for fixation were seeded in 24 well plates to facilitate collection. Rings are washed with pre-warmed PBS, followed by 30-minute fixation at room temperature with 4% Formaldehyde EM-Grade (Electron Microscopy Science #15710). Samples are collected in a conical tube and centrifuged at 2000g for 10 minutes at 4°C. Pellets are washed with PBS followed by a second spin. After discarding the supernatant, cells are mixed in 10 *µ*l of HistoGel (ThermoScientific #HG-40000-012). The mixture is shortly incubated on ice for 5 minutes to solidify the pellets before transferring to a histology cassette for standard embedding and sectioning. The slides are baked at 45°C for 20 minutes and de-paraffinized in xylene followed by washes in ethanol and D.I. water. Endogenous peroxidases are blocked with Peroxidazed-1 (Biocare Medical #PX968M) at RT for 5 minutes. Antigen retrieval is performed in a NxGEN Deloaking Chamber (Biocare Medical) using Diva Decloacker (Biocare Medical #DV2004LX) at 110°C for 15 minutes for Ki-67/Caspase-3, PAX8 and p53 (Biocare Medical #PPM240DSAA) staining or using Borg Decloacker (Biocare Medical #BD1000 S-250) at 90°C for 15 minutes for Anti-P Glycoprotein (Abcam #EPR10364-57) staining. For EGFR staining, antigen retrieval is perfomed enzymatically with Carezyme III Pronase (Biocare Medical #PRT957) at 37°C for 5 minutes.

Blocking is performed at RT for 30 minutes with Background Punisher (Biocare Medical #BP947H) at RT for 15 minutes for the EGFR staining. Primary antibodies are diluted in Da Vinci Green Diluent (Biocare Medical #PD900L) for Anti-P Glycoprotein (1:300), p53 (1:200, Biocare Medical #CME298A) and PAX8 (1:1000, Proteintech #10336-1-A) incubated at 4°C overnight or Van Gogh Diluent (Biocare #PD902H) for EGFR (1:30) incubated at RT for 30 minutes. The combo Ki-67/Caspase-3 solution is pre-diluted and added to the sample for 60 minutes at room temperature. Secondary antibody staining is performed with Rabbit on Rodent HRP-polymer (Biocare Medical #RMR622G) for the Anti-P Glycoprotein, p53 and PAX8 staining or with Mouse on Mouse HRP-polymer (Biocare Medical #MM620G) for EGFR. MACH 2 double Stain 2 (Biocare Medical #MRCT525G) is used for Ki-67/Caspase-3 combinatorial staining. All secondary antibodies are incubated at RT for 30 minutes.

Chromogen development is performed with Betazoid DAB kit (Biocare Medical #BDB2004) for Anti-P Glycoprotein, pTEN and EGFR and Ki-67 or Warp Red Chromogen Kit (Biocare Medical #WR806) for Caspase-3. The reaction is quenched by dipping the slides in D.I water. Hematoxylin-1 (Thermo Scientific #7221) is used for counterstaining. The slides are mounted with Permount (Fisher Scientific #SP15-100). Images are acquired with a Revolve Upright and Inverted Microscope System (Echo Laboratories).

## SUPPLEMENTARY MATERIAL

**Table S1.** Characteristics of samples included in this study.

Stable Lines:
| Specimen | Tumor Classification | Base Medium |
| --- | --- | --- |
| **MCF7** | Invasive breast ductal carcinoma | RPMI |
| **MD-MBA-468** | Breast adenocarcinoma | RPMI |
| **PANC1** | Pancreatic ductal adenocarcinoma | DMEM |
| **PANC03.27** | Pancreatic adenocarcinoma | RPMI |
| **HUPT4** | Pancreatic adenocarcinoma | DMEM |
| **PC3** | Prostatic adenocarcinoma | DMEM |
| **DU145** | Prostatic carcinoma | DMEM |
| **SK-NEP-1** | Ewing sarcoma | McCoy |

Clinical Specimens:
| Specimen | Tumor Classification | Stage | Sample Type | Therapy |
| --- | --- | --- | --- | --- |
| **Patient #1** | High-grade mixed type carcinoma with a high grade serous (40%) and a clear cell carcinoma (60%) component | IV | Ascites | None |
| **Patient #2** | High grade serous ovarian carcinoma | IVB | Ascites | Carboplatin / Taxol / Avastin |
| **Patient #3** | Carcinosarcoma of the ovary with malignant mixed Mullerian tumor | IIIC | Tumor | Carboplatin / Taxol |
| **Patient #4** | High grade serous carcinoma, primary peritoneal carcinoma | IIIC | Tumor | None |

**Figure S1.**
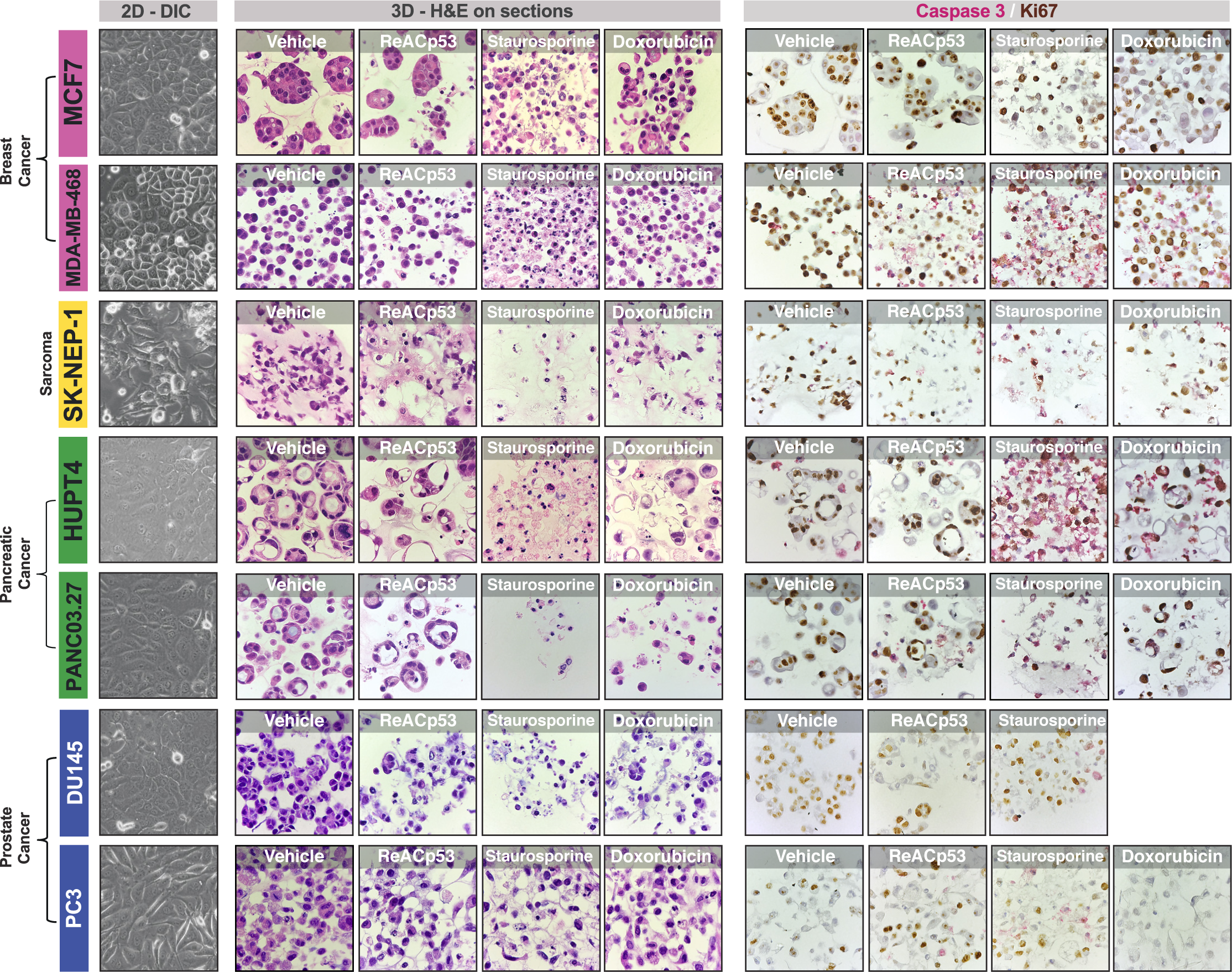
Histology of 3D tumor models. Tumor cell lines used in this study grown in 3D processed for histology. The corresponding cells grown in 2D are shown on the left (40x magnification). On the right, H&E and Caspase/Ki67 staining on sections from embedded 3D tumor organoid samples (60x magnification).

**Figure S2.**
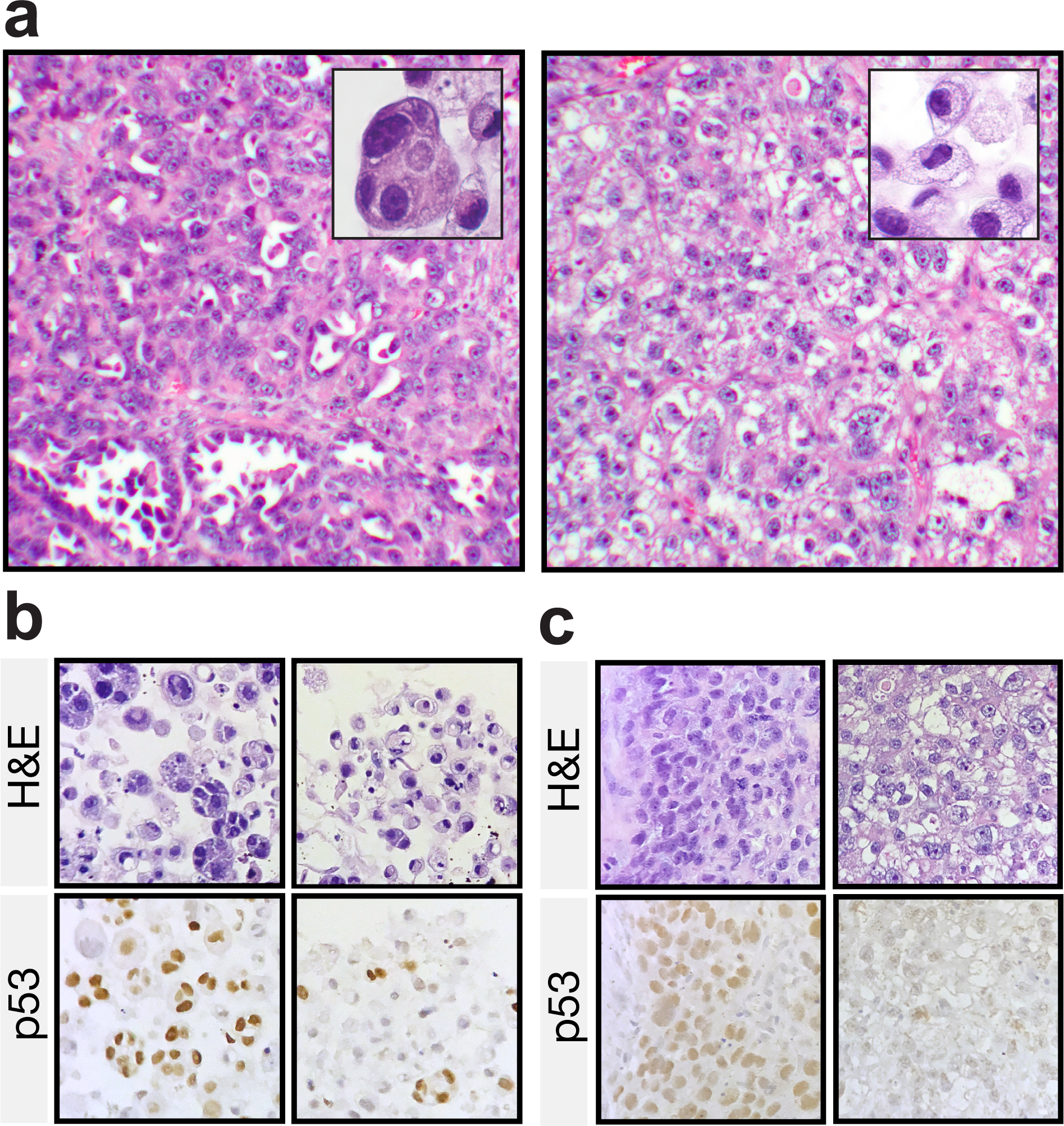
Tissue sections of high-grade mixed carcinoma from Patient #1. **(a)** Different areas of the same tumor show distinct morphologies, with a high-grade serous component on the left and clear cell carcinoma aspects on the right. 10x magnification. The insets show a representative area of the tumor orgnanoids derived from the same sample. **(b)** p53 staining of Patient #1 organoids. The H&E image is representative of the overall area, but due to the limited thickness of the sample does not show the same exact organoids of the p53 staining. **(c)** p53 staining of Patient #1 primary tumor. In both **(b)** and **(c)** two populations of cells, a p53-positive (serous) and a p53-negative (clear cell), is evident.

**Figure S3.**
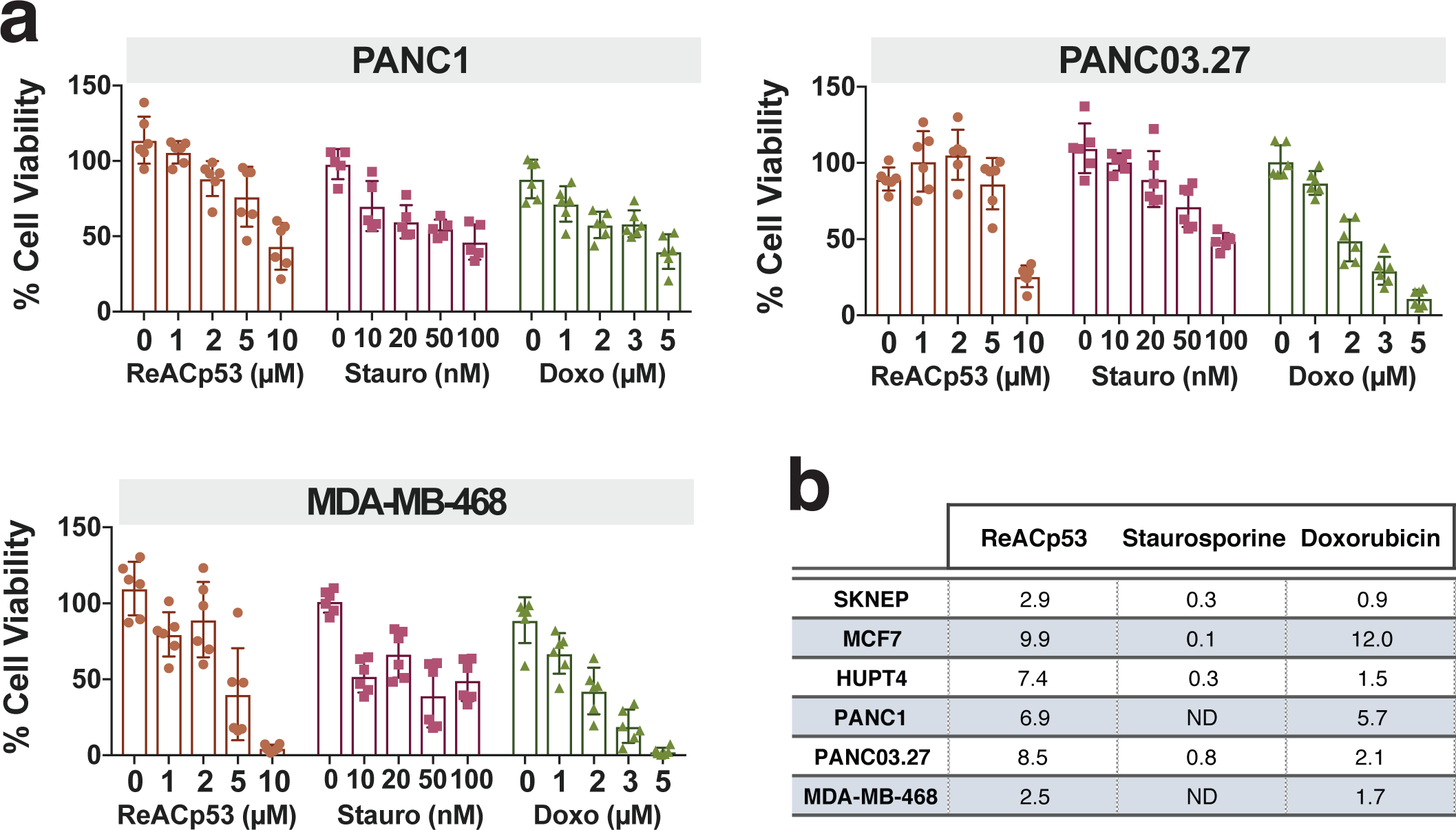
ATP readout and EC_50_ values for three-drug assay. **(a)** ATP quantification as measured by CellTiter-Glo 3D. Data from 2 independent experiments, n=3 for each are plotted. Error bars represent standard deviation; bars represent mean values. **(b)** EC_50_ values as calculate from the ATP quantification data. All values are expressed in *µ*M.

**Figure S4.**
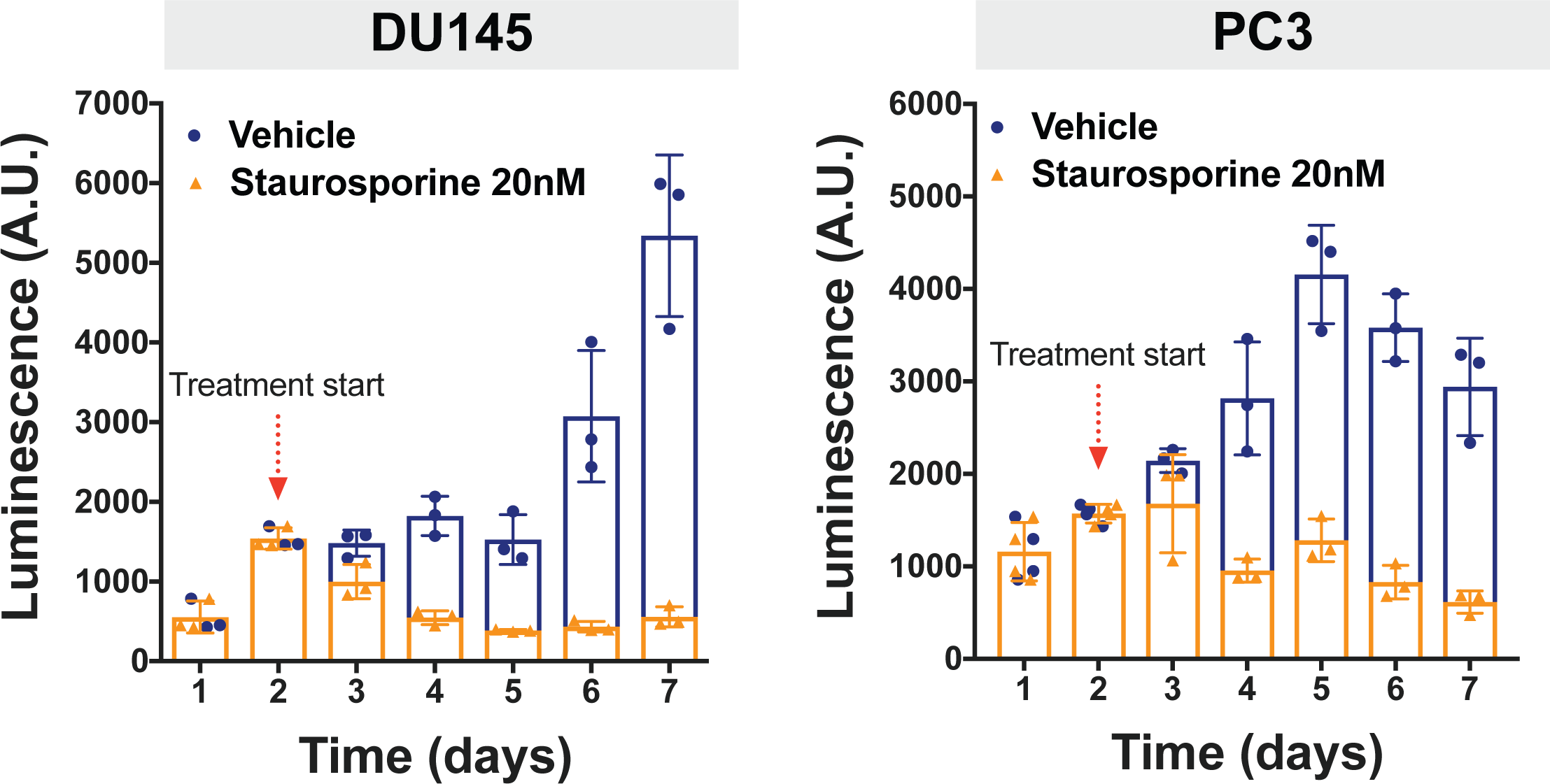
Adaptability of miniring assay to different treatment schedules. ATP quantification as measured by CellTiter-Glo 3D of prostate cancer organoids treated for 5 consecutive days with either vehicle or 20 nM Staurosporine.

**Figure S5.**
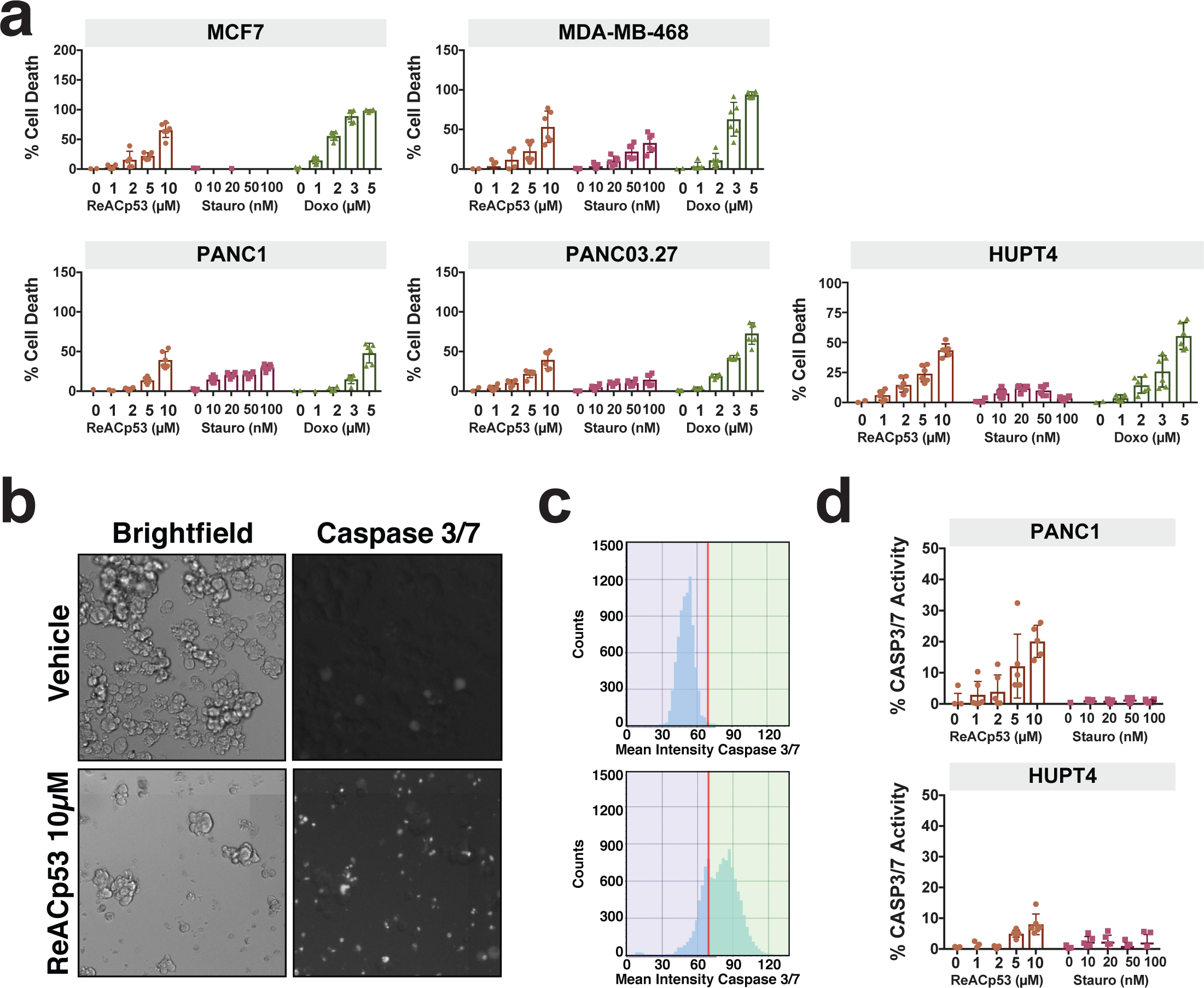
Additional optimized readouts for mini-ring assay. **(a)** Quantification of the calcein release / PI uptake experiment. Two independent experiments shown, n=3 for each. Error bars are standard deviation while bars represent mean values. **(b)** and **(c)** Example of outcome for the caspase 3/7 cleavage experiment. DU145 prostate cancer cells are shown. A substrate becomes fluorescent when cleaved by caspase 3 or 7. Treatment induces high levels of caspase activation. Histograms of fluorescence intensity are shown in **(c)**. **(d)** Quantification of active caspase 3/7 activity normalized to control. Doxorubicin has intrinsic fluorescence that masks the caspase signal hence was excluded from this analysis.

**Figure S6.**
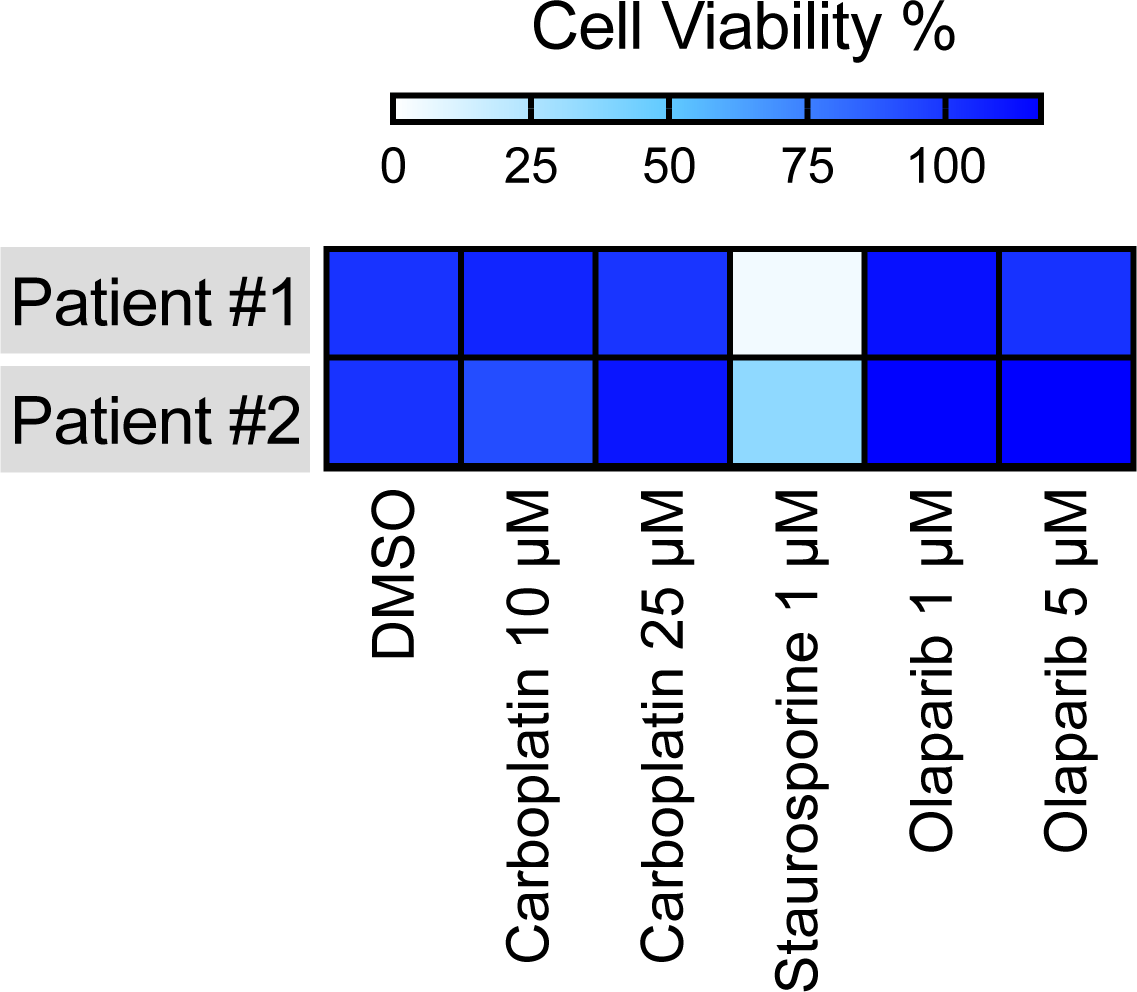
Response of PDTOs to chemotherapy. PDTOs established from Patients #1 and #2 were also tested for response to Carboplatin and Olaparib.

**Figure S7.**
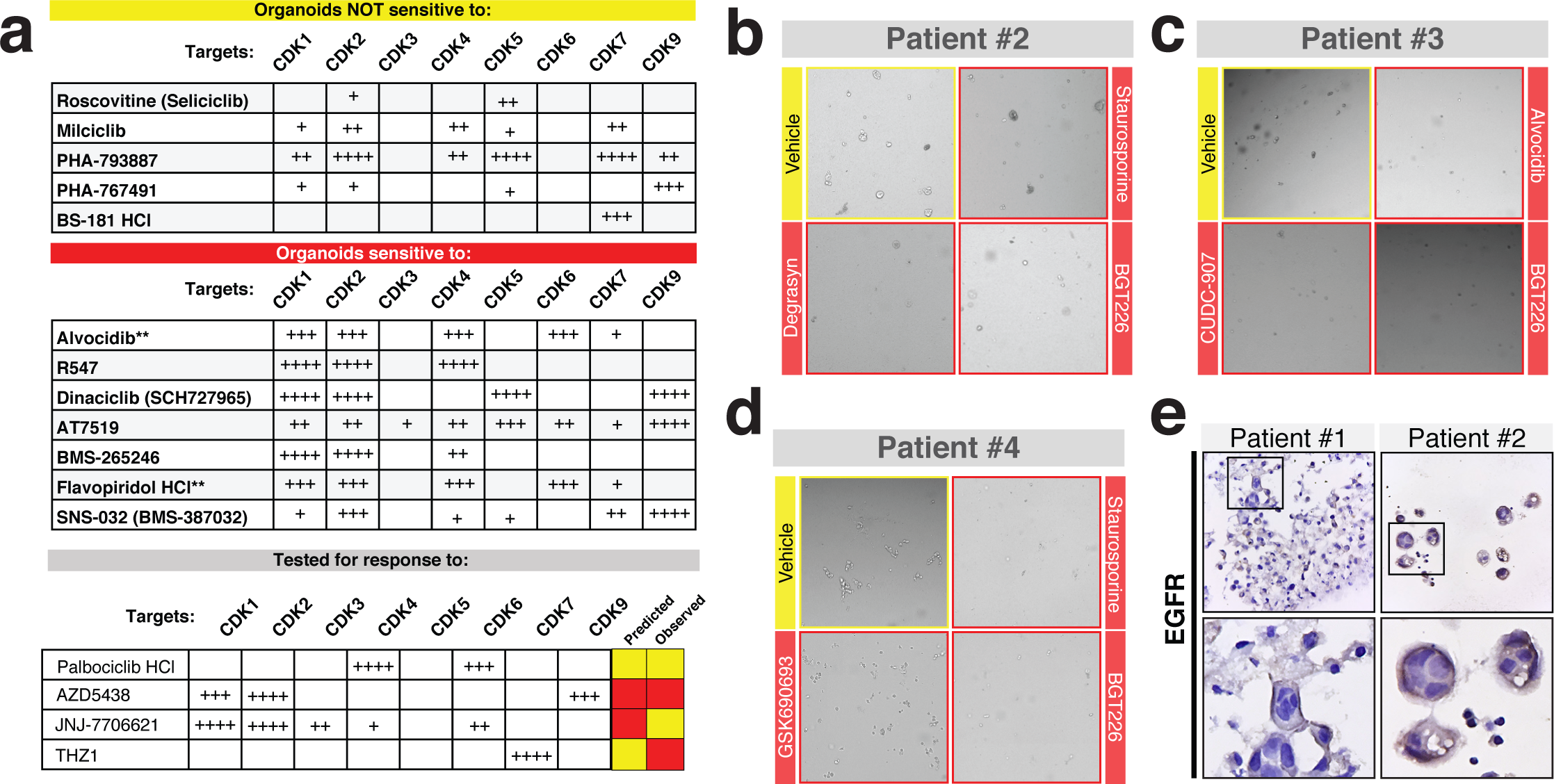
Results and validation of PDTO kinase screening. **(a)** List of CDK inhibitors that did or did not induce cell death in >75% Patient #1’s organoids. Targets and specificity of each is listed. The patient responded to CDK inhibitors hitting CDK1/2 in combination with CDK 4/6 or CDK 5/9. **(b), (c) and (d)** Representative images of post-treatment and post-dispase PDTOs. **(e)** Expression of EGFR in Patient #1 and Patient #2 3D tumors. Magnification: 40x.

## Notes

The authors declare no potential conflict of interest.

## References

1. Cummings, C. A., Peters, E., Lacroix, L., Andre, F. & Lackner, M. R. The Role of Next-Generation Sequencing in Enabling Personalized Oncology Therapy. Clin. Transl. Sci. 9, 283–292 (2016).

2. Simon, R. & Roychowdhury, S. Implementing personalized cancer genomics in clinical trials. Nat. Rev. Drug Discov. 12, 358–369 (2013).

3. Letai, A. Functional precision cancer medicine-moving beyond pure genomics. Nat. Med. 23, 1028–1035 (2017).

4. Vlachogiannis, G. et al. Patient-derived organoids model treatment response of metastatic gastrointestinal cancers. Science 359, 920–926 (2018).

5. Voest, E. E. & Bernards, R. DNA-Guided Precision Medicine for Cancer: A Case of Irrational Exuberance? Cancer Discov. 6, 130–132 (2016).

6. Prasad, V., Fojo, T. & Brada, M. Precision oncology: origins, optimism, and potential. Lancet Oncol. 17, e81–e86 (2016).

7. Tannock, I. F. & Hickman, J. A. Limits to Personalized Cancer Medicine. N. Engl. J. Med. 375, 1289–1294 (2016).

8. Pickl, M. & Ries, C. H. Comparison of 3D and 2D tumor models reveals enhanced HER2 activation in 3D associated with an increased response to trastuzumab. Oncogene 28, 461–468 (2009).

9. Katt, M. E., Placone, A. L., Wong, A. D., Xu, Z. S. & Searson, P. C. In Vitro Tumor Models: Advantages, Disadvantages, Variables, and Selecting the Right Platform. Front. Bioeng. Biotechnol. 4, 12 (2016).

10. Tanner, K. & Gottesman, M. M. Beyond 3D culture models of cancer. Sci. Transl. Med. 7, 283ps9 (2015).

11. Nyga, A., Cheema, U. & Loizidou, M. 3D tumour models: novel in vitro approaches to cancer studies. J. Cell Commun. Signal. 5, 239–248 (2011).

12. Fong, S., Debs, R. J. & Desprez, P.-Y. Id genes and proteins as promising targets in cancer therapy. Trends Mol. Med. 10, 387–392 (2004).

13. Kimlin, L. C., Casagrande, G. & Virador, V. M. In vitro three-dimensional (3D) models in cancer research: an update. Mol. Carcinog. 52, 167–182 (2013).

14. Pauli, C. et al. Personalized In Vitro and In Vivo Cancer Models to Guide Precision Medicine. Cancer Discov. 7, 462–477 (2017).

15. Halfter, K. & Mayer, B. Bringing 3D tumor models to the clinic - predictive value for personalized medicine. Biotechnol. J. 12, (2017).

16. Soragni, A. et al. A Designed Inhibitor of p53 Aggregation Rescues p53 Tumor Suppression in Ovarian Carcinomas. Cancer Cell 29, 90–103 (2016).

17. Breslin, S. & O’Driscoll, L. Three-dimensional cell culture: the missing link in drug discovery. Drug Discov. Today 18, 240–249 (2013).

18. Breslin, S. & O’Driscoll, L. The relevance of using 3D cell cultures, in addition to 2D monolayer cultures, when evaluating breast cancer drug sensitivity and resistance. Oncotarget 7, 45745–45756 (2016).

19. Friedrich, J., Seidel, C., Ebner, R. & Kunz-Schughart, L. A. Spheroid-based drug screen: considerations and practical approach. Nat. Protoc. 4, 309–324 (2009).

20. Zanoni, M. et al. 3D tumor spheroid models for in vitro therapeutic screening: a systematic approach to enhance the biological relevance of data obtained. Sci. Rep. 6, 19103 (2016).

21. Kelm, J. M., Timmins, N. E., Brown, C. J., Fussenegger, M. & Nielsen, L. K. Method for generation of homogeneous multicellular tumor spheroids applicable to a wide variety of cell types. Biotechnol. Bioeng. 83, 173–180 (2003).

22. Boj, S. F. et al. Organoid models of human and mouse ductal pancreatic cancer. Cell 160, 324–338 (2015).

23. Walsh, A. J. et al. Quantitative optical imaging of primary tumor organoid metabolism predicts drug response in breast cancer. Cancer Res. 74, 5184–5194 (2014).

24. Francies, H. E., Barthorpe, A., McLaren-Douglas, A., Barendt, W. J. & Garnett, M. J. Drug Sensitivity Assays of Human Cancer Organoid Cultures. Methods Mol. Biol. Clifton NJ (2016). doi: 10.1007/7651_2016_10

25. Belmokhtar, C. A., Hillion, J. & Ségal-Bendirdjian, E. Staurosporine induces apoptosis through both caspase-dependent and caspase-independent mechanisms. Oncogene 20, 3354–3362 (2001).

26. Lovitt, C. J., Shelper, T. B. & Avery, V. M. Doxorubicin resistance in breast cancer cells is mediated by extracellular matrix proteins. BMC Cancer 18, 41 (2018).

27. Zhou, Q. Targeting Cyclin-Dependent Kinases in Ovarian Cancer. Cancer Invest. 35, 367–376 (2017).

28. Li, B. et al. Therapeutic Rationale to Target Highly Expressed CDK7 Conferring Poor Outcomes in Triple-Negative Breast Cancer. Cancer Res. 77, 3834–3845 (2017).

29. Ben-David, U. et al. Patient-derived xenografts undergo mouse-specific tumor evolution. Nat. Genet. 49, 1567–1575 (2017).

30. Eirew, P. et al. Dynamics of genomic clones in breast cancer patient xenografts at single-cell resolution. Nature 518, 422–426 (2015).

31. Sun, S. et al. Prognostic Value and Implication for Chemotherapy Treatment of ABCB1 in Epithelial Ovarian Cancer: A Meta-Analysis. PLOS ONE 11, e0166058 (2016).

32. Vaidyanathan, A. et al. ABCB1 (MDR1) induction defines a common resistance mechanism in paclitaxel- and olaparib-resistant ovarian cancer cells. Br. J. Cancer 115, 431–441 (2016).

33. Hirte, H. et al. A phase II study of erlotinib (OSI-774) given in combination with carboplatin in patients with recurrent epithelial ovarian cancer (NCIC CTG IND.149). Gynecol. Oncol. 118, 308–312 (2010).

34. Rauh-Hain, J. A., Birrer, M. & del Carmen, M. G. “Carcinosarcoma of the ovary, fallopian tube, and peritoneum: Prognostic factors and treatment modalities”. Gynecol. Oncol. 142, 248–254 (2016).

35. Mano, M. S. et al. Current management of ovarian carcinosarcoma. Int. J. Gynecol. Cancer 17, 316–324 (2007).

36. Rhodes, N. et al. Characterization of an Akt Kinase Inhibitor with Potent Pharmacodynamic and Antitumor Activity. Cancer Res. 68, 2366–2374 (2008).

37. Markman, B. et al. Phase I safety, pharmacokinetic, and pharmacodynamic study of the oral phosphatidylinositol-3-kinase and mTOR inhibitor BGT226 in patients with advanced solid tumors. Ann. Oncol. Off. J. Eur. Soc. Med. Oncol. 23, 2399–2408 (2012).

38. Massacesi, C. et al. PI3K inhibitors as new cancer therapeutics: implications for clinical trial design. OncoTargets Ther. 9, 203–210 (2016).

39. Markman, B., Atzori, F., Pérez-García, J., Tabernero, J. & Baselga, J. Status of PI3K inhibition and biomarker development in cancer therapeutics. Ann. Oncol. 21, 683–691 (2010).

40. Dumbrava, E. I., Meric-Bernstam, F. & Yap, T. A. Challenges with biomarkers in cancer drug discovery and development. Expert Opin. Drug Discov. 13, 685–690 (2018).

41. Huang, L. et al. Ductal pancreatic cancer modeling and drug screening using human pluripotent stem cell-and patient-derived tumor organoids. Nat. Med. 21, 1364–1371 (2015).

42. Ben-David, U. et al. Genetic and transcriptional evolution alters cancer cell line drug response. Nature 1 (2018). doi: 10.1038/s41586-018-0409-3

43. Gock, M. et al. Tumor Take Rate Optimization for Colorectal Carcinoma Patient-Derived Xenograft Models. BioMed Res. Int. 2016, (2016).

44. Tate Thigpen, J., Blessing, J. A., DeGeest, K., Look, K. Y. & Homesley, H. D. Cisplatin as initial chemotherapy in ovarian carcinosarcomas: a Gynecologic Oncology Group study. Gynecol. Oncol. 93, 336–339 (2004).

